# The Body Mirroring Thought: The Relationship Between Thought Transitions and Fluctuations in Autonomic Nervous Activity Mediated by Interoception

**DOI:** 10.1101/2024.07.30.605929

**Authors:** Mai Sakuragi, Kazushi Shinagawa, Yuri Terasawa, Satoshi Umeda

**Affiliations:** Department of Psychology, Keio University, 2-15-45 Mita, Minato-ku, Tokyo, 108-8345, Japan; Japan Society for the Promotion of Science, Kojimachi Business Center Building, 5-3-1, Kojimachi, Chiyoda-ku, Tokyo 102-0083, Japan; Keio University Global Research Institute, 2-15-45 Mita, Minato-ku, Tokyo, 108-8345, Japan

**Keywords:** Mind Wandering, Autonomic Nervous Activity, Interoception

## Abstract

Our thought states change unconsciously. This study verified that the transference of thought states varies with fluctuations in autonomic nervous activity, and that this effect is modulated by interoceptive accuracy. The participants completed the heartbeat counting task (HCT) and vigilance task (VT). We assessed the participants’ interoceptive accuracy based on their performance on the HCT. The VT is a simple attention task, and during this task, we asked the participants to report the content and contemplation of their thoughts. Consequently, participants with accurate interoception were more likely to remain in a highly contemplative thought state when sympathetic activity was activated. In contrast, the dominance of parasympathetic activity facilitated transitions to different thought states or experiences of less contemplative thought states in them. The results suggest that even subtle changes in bodily responses at rest can affect thought transitions in people with accurate interoception.

## 1. Introduction

Our thoughts unconsciously shift to different states. The phenomenon of thoughts wandering to matters unrelated to a task has been studied under the name of mind-wandering (MW; Smallwood & Schooler, 2015). MW research in experimental psychology and cognitive neuroscience is conducted by presenting participants with thought probes in which they respond to their previous thoughts (probe-caught method; Weinstein, 2018; Weinstein et al., 2018) or by having them spontaneously report their thought deviations (self-caught method; Chu et al., 2023) while engaging in simple tasks with a low attentional load. Although MW studies have increased since the early 2000s, its definitions and measurement methods still need to be standardized. Most early MW studies focused on the content of thought. For example, MW was defined as task-unrelated thought, and task-related/unrelated dichotomies or Likert scale have been widely used to evaluate the task-relatedness of participants’ thoughts (Seli et al., 2018). On the other hand, some measures divide the content of thoughts into several categories based on dimensions such as task relevance and stimulus independence (whether the content of thoughts is related to stimuli or environmental changes presented during the task). Participants select the category that is most appropriate for the content of their thoughts (Stawarczyk et al., 2011). Based on these previous researches, using the Likert scale or Visual Analog Scale to create a gradation of task relevance or to have participants select the appropriate thought content from multiple categories rather than a task-related/unrelated dichotomy is more appropriate (Kane et al., 2021; Seli et al., 2018).

In recent years, components other than thought content, such as contemplation (subjective concentration on a specific thought content), intention, and intrusiveness, have become subjects of research (Christoff et al., 2016; Jubera-García et al., 2020; Kane et al., 2021; Seli et al., 2016; Smallwood et al., 2016). These studies revealed that different contemplation and intentionality of thoughts affected not only the participants’ subjective states of consciousness, but also their performance on parallel attention tasks and physiological measures, such as pupil dilation. These findings suggest that different types of thoughts may have been mixed and grouped as task-related or task-unrelated in previous studies. In addition, a dynamic model that focuses on temporal variation rather than individual thoughts has been proposed (Christoff et al., 2016; Mittner et al., 2016). Studies analyzing temporal changes in thought content also indicate complex states in which multiple thought contents coexist (Shinagawa et al., 2023; Zanesco et al., 2020). In light of these studies, to understand our thoughts, it is necessary to estimate the state of thought, which integrates multiple components of thought, and to address its temporal variation.

Transitions in thought states may be related to fluctuations in autonomic nervous activity. This system regulates critical physiological functions, including the cardiac, respiratory, digestive, vasomotor, and endocrine systems. Ottaviani and her colleagues defined negative and contemplative thoughts, such as rumination and worrying, as perseverative cognition (Ottaviani et al., 2013; Ottaviani, Shahabi, et al., 2015). They reported that perseverative cognition differs from MW in the variability of autonomic nervous activity, as estimated from heart rate variability (HRV) and blood pressure. For example, when participants’ thought states were measured using the probe-caught method during a simple attention task, perseverative cognition was accompanied by a higher heart rate and lower RMSSD (parasympathetic index), one of the HRV indices, compared to MW (Ottaviani et al., 2013). This phenomenon has also been observed in patients with perseverative cognition as a clinical symptom, such as major depression (Ottaviani, Medea, et al., 2015; Ottaviani, Shahabi, et al., 2015) and generalized anxiety disorder (Ottaviani et al., 2016). These results suggest that the perseverance of thought content influences the degree of variability in autonomic nervous activity. However, the psychological and neural processes linking thought states and autonomic nervous activity have yet to be demonstrated experimentally.

As mentioned in the previous paragraphs, fluctuations in autonomic nervous activity result in changes in bodily responses, such as cardiac activity and respiration. Interoception is the perception and information processing associated with changes in bodily responses. It refers to the entire process by which the nervous system perceives, interprets, and integrates signals transmitted from within the body, particularly from internal organs, and operates across both conscious and unconscious levels (Khalsa et al., 2018). Given that the subjective trigger is unknown for more than half of the MW reported in daily life (Faber & D’Mello, 2018), the perception of a change in physical state at an unconscious level may influence thought shifting. Many studies of interoception have focused on insular cortex, which is the central hub of interoception. This region, particularly the anterior part of it, mediates the dynamic interaction of two brain functional networks: the default mode network (DMN), which is involved in MW, and the central executive network, which is involved in the active maintenance and manipulation of information in working memory (Menon, 2011; Menon & Uddin, 2010; Uddin, 2015). In light of these findings, fluctuations in autonomic nervous activity and the input of updated interoceptive information to the insular cortex may cause switching in neural activity, resulting in transitions in thought states. There is also a relationship between the heartbeat-evoked response (HER), an afferent signal of cardiac activity in the magnetoencephalography, one of the neural indicators of interoception, and the state of self-expression during MW (Babo-Rebelo, Richter, et al., 2016; Babo-Rebelo, Wolpert, et al., 2016). These studies revealed that the level of self-expression during MW and the HER amplitude in the DMN covaried. Although these studies found no direct relationship between peripheral autonomic nervous activity and thought, they suggest that the neural processing of unconscious bodily responses may contribute to generating specific thought states.

Based on the findings described above, the present study had two objectives: first, to estimate the state of thought by integrating the thought content and the contemplation of how deeply the participant was thinking about that content; second, to examine the relationship between the transition of estimated thought states and fluctuations in autonomic nervous activity and interoception. Participants performed the vigilance task (VT), a simple attention task used in the laboratory to measure spontaneous transitions in thought states (Kvavilashvili et al., 2020). Participants responded by pressing a keyboard only when the target stimuli were presented. During this task, they answered the characteristics of the immediately preceding thought in response to randomly presented thought probes. Transition series were estimated using hidden Markov models (HMM) to reveal patterns of change in thought content and contemplation over time. Autonomic nervous activity was estimated using an HRV analysis of pulse data. Additionally, the present study focused on interoceptive accuracy (objective accuracy in detecting bodily states), a subcategory of interoception (Garfinkel & Critchley, 2013). Participants completed the heartbeat counting task (HCT) to measure this index (Schandry, 1981). Individuals with higher performance on this task have stronger functional connectivity in areas associated with interoception (Chong et al., 2017) and greater amplitude of afferent signals associated with cardiac activity (Coll et al., 2021) in the resting state. This suggests that participants with higher interoceptive accuracy, as measured by the HCT, may have enhanced neural processing of interoception, even when their attention is not focused on a bodily response. We used a linear mixed model to estimate the relationship between the transition probabilities and transition patterns of thought states, autonomic nervous activity during the task, and interoceptive accuracy. DMN, which is activated during MW, is also activated when participants are at rest, not engaged in a specific task (Greicius et al., 2003, 2009; Raichle, 2015). From this, we formulated the following two hypotheses: (1) when parasympathetic activity is dominant and the body is relaxed, thought state transitions are more frequent, and less contemplative thought states are more likely to occur; conversely, when sympathetic activity is activated, thought state transitions are less frequent and highly contemplative thought states are more likely to occur, and (2) the more accurate the interoception, the more likely it is that information processing about these changes in autonomic nervous activity is facilitated and that transitions in thought states are more likely to occur.

## 2. Material and Methods

### 2.1. Participants

One hundred university and graduate students (70 women, 30 men, *M_age_* = 21 ± 1.46 years) were recruited. All participants had normal or corrected-to-normal vision. This study was approved by the Keio University Research Ethics Committee (no. 210300000) and was conducted in accordance with the Declaration of Helsinki. All participants provided written informed consent before participation. As indicated in a previous paper, the primary objective of the present study was to examine how the transition patterns of thought changed during the manipulation of heart rate by presenting subthreshold vibration stimuli (Sakuragi et al., 2023). The presentation of subthreshold vibration stimuli during the task was not explained in the pre-experiment because of the possibility of influencing the heart rate change effect of the stimuli. After the experiment was completed, the presentation of subliminal vibration during the VT and its purpose were disclosed, and written informed consent was again obtained for using the data obtained in the experiment. One participant was excluded from the analysis because the participant responded to the same category in all 20 responses to the thought probe, which precluded HMM parameter estimation (section 2.5.1). In addition, three participants whose number of responses to mind-blanking (thinking about nothing) exceeded the overall mean +2 standard deviations (SD) were also excluded from the analysis because their reports on the thought state data were deemed unreliable, as they were significantly more likely to have a decreased level of consciousness compared to the other participants.

### 2.2. Apparatus

We used a pulse oximeter (NONIN, 8600) and a cuff (NONIN, 8000SM) attached to the left index finger’s tip to measure participants’ pulse rates. The measured signal was recorded by POWER1401mk2 and Spike2 (Cambridge Electronic Design). We used two cushions as armrests (Cushion L and Cushion R). Participants placed their left forearm on Cushion L and their right elbow on the left half of Cushion R.

### 2.3. Procedure

Participants came to the laboratory and performed two tasks: the HCT and the VT. Participants put a pulse oximeter on their left index finger in both tasks.

#### 2.3.1. The Heartbeat Counting Task (HCT)

The participants first completed the HCT to measure their interoceptive accuracy (Schandry, 1981). This task comprised three processes: resting heart rate measurement, the HCT, and the time estimation task (TET). While performing these tasks, the participants placed their entire left arm on Cushion L on the desk. In the HCT, they silently counted their heartbeats over time intervals (2 × 25 s, 2 × 35 s, and 2 × 45 s) and then reported the number of heartbeats counted for each interval. They were instructed not to count their heart rate by touching themselves, pressing their bodies hard against a backrest or desk, or holding their breath. In addition, referring to an adapted version of the teaching of the HCT (Desmedt et al., 2018), participants were asked not to guess and count heartbeats they did not feel. Lastly, they estimated the length of the time they felt had elapsed at a series of time intervals (2 × 23 s, 2 × 49 s, and 2 × 56 s) and then reported that length (TET). In the HCT and TET, each interval was measured in random order.

#### 2.3.2. The Vigilance Task (VT)

After the HCT, we placed Cushion R on the desk, and the participants rested their right elbows there. The participants wore silicone earplugs in both ears. Then, we started the second task, the VT. Participants performed a simple attention task for a sustained period, and the shifting of thought states during the task was measured. At first, the cross of the gazing point was presented for 1 s. After that, various images of straight lines were presented. We prepared two types of stimuli for the VT: the rows of images with multiple straight lines aligned horizontally (control stimulus) and the row of images with four straight lines aligned vertically (target stimulus). The presentation time of one image and the time until one image disappeared and the next image appeared were 500 ms each. The participants responded by pressing the space key as soon as the target stimuli appeared on the screen. Contrastingly, we instructed participants to stare only at the screen when the control stimuli were presented. The participants’ responses were recorded for the 1 s between the presentation of one image and the next. In one trial, we presented 42 images (39 × control stimuli and 3 × target stimuli). Target stimuli were presented at random times per trial. The tasks were performed for 20 trials.

On each trial, three thought probes were presented after all the image stimuli were presented: thought content, contemplation, and consistency. This frequency of presenting probes is generally consistent with the average duration of MW self-caught in previous studies (Voss et al., 2018). These thought probes were presented 20 times per block, based on the previous studies that measured the content of thoughts and their shifting by probe-caught, as in this study (Kane et al., 2016, 2017; Welhaf et al., 2020; Zanesco et al., 2020).

In thought probe (1), participants reported their contents of thoughts just before the probe was displayed. The present study categorized thought content into task relevance and stimulus independence (Stawarczyk et al., 2011). We prepared the following six categories for participants to categorize their thoughts: 1) task-focused; 2) task-related (the score of the task/elapsed time); 3) exteroception (sound/flash); 4) interoception (hunger/pain/heartbeat); 5) self-related (past episodes, plans for the future); and 6) mind-blanking (nothing). In our previous study, when we used categories classified according to the same criteria, each category was selected around five times out of 20, except for 3) exteroception, where the input was restricted by stimulus control and earplugs in the experiment (Sakuragi et al., 2023; Supplementary data). This suggests that the participants could correctly evaluate their thought contents according to these categories. Since the purpose of this study was to discuss the role of interoception in the transition of thought states, as distinguished from exteroception (i.e., the perception of peripheral stimuli), we separated Categories 3 and 4. Additionally, we gave past episodes and future plans about oneself as specific examples of Category 5. This is based on previous research showing that during MW, various self-referential thoughts are reported, including simulations of one’s past episodes and future plans and goals (D’Argembeau, 2018; Kvavilashvili et al., 2020). Mind-blanking refers to a state in which participants could not report their thought state because they were unaware of any internal or external stimuli (Kaufmann et al., 2024; Ward & Wegner, 2013).

When participants pressed one of the number keys from 1 to 5 in thought probe (1), thought probe (2) appeared. However, for participants who answered 6 in thought probe (1), the following probes (2) and (3) were not presented. In thought probe (2), participants responded by selecting one of the options from “1. not very much,” “2. deeply,” and “3. very deeply” to indicate the depth they contemplated the thoughts they responded to in probe (1). When participants pressed one of the number keys from 1 to 3 in probe (2), thought probe (3) appeared. Participants responded by pressing the number key: 1 if they were thinking about a single matter; 2 if they were thinking about multiple contents and their semantic relevance was high; or 3 if their semantic relevance was low. When participants pressed one of the number keys 1‒3 in probe (3), the gazing point was presented for 1 s, and the presentation of the linear image was resumed. Only data from probes (1) and (2) were used in this study.

### 2.4. Pre-Processing

#### 2.4.1. The HCT and the TET

We performed pre-processing on the first task, the HCT and TET error rates. We calculated the error rates of the HCT using the following formula:

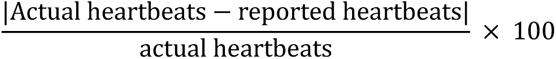

Six error rates were obtained for each participant and those values were averaged to obtain the individual’s HCT error rate. We calculated the error rates for the TET using the following formula:

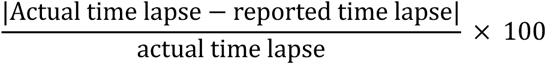

Six error rates were obtained for each participant, and those values were averaged to obtain the individual’s TET error rate. To confirm that the HCT results were not affected by inferences based on time estimates, we calculated Pearson’s product-moment correlation coefficient for the HCT and the TET error rates.

#### 2.4.2. Heart Rate Variability (HRV)

We performed HRV analysis to index fluctuations in participants’ autonomic nervous activity during the task. In this study, we calculated three indices using R (version 4.3.2): LF/HF, HF, and RMSSD. LF/HF reflects the relative balance between the sympathetic and parasympathetic nervous activity, while HF and RMSSD reflect parasympathetic activity (Ernst, 2017). LF/HF and HF were used to quantify autonomic nervous activity throughout the task, which lasted less than 20 minutes, and to examine their relationship to thought transition patterns. RMSSD was used to examine the relationship between the thought content and autonomic nervous activity on a trial-by-trial basis. RMSSD is a time-domain index that quantifies autonomic nervous activity over a relatively short period (Nussinovitch et al., 2011). However, because it is a simple index calculated with a small amount of data, the robustness of its evaluation may be insufficient. Therefore, in this study, we also used LF/HF and HF, which are frequency-domain indices, to examine whether similar trends can be observed throughout the entire task and on a trial-by-trial basis. In calculating these indices, the pulse wave interval, the interval between the sharpest portions of the pulse wave, was resampled to 100 milliseconds. HRV calculated using the pulse interval is sufficiently accurate when the participant is at rest (Schäfer & Vagedes, 2013). Additionally, as in the previous study (Ottaviani et al., 2013), we excluded pulse data while participants responded to the thought probe and more than 30 seconds before each thought probe presentation. The second exclusion was made to remove the influence of nervousness immediately after the trial’s start. LF/HF and HF were calculated by performing a spectral analysis using the fast Fourier transform algorithm on the time series of pulse interval data obtained from the entire task. The low-frequency component at 0.04-0.15 Hz was defined as LF, and the high-frequency component at 0.15-0.5 Hz as HF, and LF/HF was calculated from the two values. RMSSD was calculated as follows: We first squared the differences between successive pulse rates for each trial. Then, we averaged these squared differences. Finally, we took the square root of this average to obtain the RMSSD for that trial. Based on the actual data distribution, four participants, one male and three females, whose LF/HF or HF values deviated from the overall mean ±3 SD, were excluded from the analysis.

### 2.5. Modeling

#### 2.5.1. Hidden Markov Model (HMM)

HMM was used to estimate the thought states generated by the content and contemplation of thoughts. This model assumes that unobservable series of states generate the observed data. Each hidden state has a probability of generating an observable output, with a specific probability of transition between states. This approach has been used in prior research to reveal the behavioral and attentional states behind the experience sampling data of the thought (Shinagawa et al., 2023; Zanesco et al., 2020) and response time for the task (Bastian & Sackur, 2013). This study used time series data of thought content and contemplation (20 probes each) as inputs (observation series) to the HMM. The HMM was implemented using the depmixS4 package in R 4.3.2 (Visser & Speekenbrink, 2010). This package uses an expectation-maximization algorithm to find maximum likelihood estimates in datasets with hidden variables, estimating HMM parameters such as state transition, output, and initial state probabilities. Based on previous studies, we determined the number of hidden states using the Bayesian Information Criterion (BIC; Shinagawa et al., 2023; Zanesco et al., 2020). BIC is a criterion measure for model selection based on a balance between model complexity (number of parameters) and fit to the data (likelihood) (Schwarz, 1978). Furthermore, to increase model stability and reproducibility, we repeated the estimation of the selected model 100 times and finally adopted the model with the lowest BIC among them. We showed each thought state’s response distributions of thought content and contemplation based on the adopted model. We also calculated the probability of occurrence for each state and the probability of transition between states. For each participant, the probability of transitions to different thought states on successive trials (transition probability) was calculated. To investigate the possibility that different autonomic nervous activity may be associated with different thought states, the mean of the RR interval and RMSSD for each state were calculated, and a one-way ANOVA was performed with the thought state as a factor. In addition, to examine whether the number of times each thought state was reported differed depending on the participant’s interoceptive accuracy, we calculated Pearson’s product-moment correlations between the participants’ scores on the HCT and the number of occurrences of each state. Because we repeated the calculation of the correlation coefficients for the four thought states, the *p*-values for the test of no correlation were corrected using the Benjamini & Hochberg method.

#### 2.5.2. Linear Regression Model

First, to clarify the relationship among the transition probability of thought states throughout the task, autonomic nervous activity, and interoceptive accuracy, we estimated linear regression models with participants’ LF/HF or HF and their HCT and TET performance as independent variables and transition probability as the dependent variable. The HF and LF/HF values were log-transformed to stabilize the model convergence. Based on the hypothesis that interoceptive accuracy mediates the effect of fluctuations in autonomic nervous activity on thought transitions, we focused only on the interaction between the indicator of autonomic nervous activity (LF/HF or HF) and interoceptive accuracy. The reason for including TET scores in the present model analysis was to exclude the influence of the accuracy of time perception on thought transitions through confounding with interoceptive accuracy (Desmedt et al., 2020). We estimated models including only LF/HF or HF and TET scores, and a model adding the main effect of the HCT score and the interaction between these autonomic nervous activity indices and the HCT score. The Widely Applicable Information Criterion (WAIC; Watanabe, 2010) was used to determine which model to adopt. This indicator estimates the expected log pointwise predictive density for a new dataset that integrates the posterior distributions of the model parameters, allowing for a combined assessment of model fit and complexity. As a lower WAIC value indicates that the model fits the observed data better, a model with a lower WAIC value was adopted.

Next, we examined the relationship between the transition patterns of thought states between the preceding and following trials, RMSSD between thought transitions, and interoceptive accuracy. For each participant, the appearance probability of each of the 16 transition patterns on consecutive trials of the four thought states and the RMSSD of the trials between each transition pattern were calculated. Generalized mixed models were constructed using RMSSD, the HCT and TET performance as independent variables, and the appearance probability of each transition pattern as the dependent variable, including individual random effects. The model included the main effects of the three independent variables and the interaction of RMSSD between the HCT scores. To stabilize the model’s convergence, all independent variable values were standardized. Based on the hypothesis that interoceptive accuracy mediates the effect of variations in autonomic nervous activity on the appearance probability of particular thought transition patterns, we focused only on the interaction effect. After estimating the models, including RMSSD, the HCT, and TET for all transition patterns, we also estimated models, removing the effect of the HCT only for those transition patterns where the interaction between RMSSD and the HCT was observed. By comparing the WAIC of the models that included the HCT and those that did not, we examined whether including the effect of the HCT in the model would allow us better to explain the appearance probability of the transition patterns.

A Bayesian approach was used to estimate the parameters of these statistical models. In repeating the model estimation for the 16 patterns of thought transitions, the posterior probabilities for each hypothesis test were computed directly to reduce the increase in the false positive rate due to multiple comparisons. We used the brms R package to analyze (Bürkner, 2017, 2018) and estimated the parameters using four Markov chain Monte Carlo chains. The thinning parameter was set to 1, and sampling was repeated 16000 times, including 2000 burn-in samples. This resulted in 14,000 posterior samples per chain, combined into a single posterior sample of 56,000 samples for each parameter. In the model estimation of the effect of HF and the HCT score on the transition probability of thought, the HF values were logarithmic transformations based on the actual data distribution. The dependent variable was assumed to follow a Gaussian distribution with a center at 0 and a scale of 5. The regression weights had Cauchy prior distributions centered at 0 with a scale of 2. The prior distributions of the intercept and residuals of the model were Student’s *t*-distributions centered at 0, with 3 degrees of freedom and a scale of 2.5, respectively. The SD of the intercept, random effects, and random effects intercepts had uninformed prior distributions. Model convergence was evaluated based on the Gelman-Rubin convergence statistic *R̂* (Gelman & Rubin, 1992), with values close to 1 indicating negligible differences in within-chain and between-chain variance. The mean and 95% confidence interval (CI) of the parameter estimates were used to examine the effect of each parameter.

## 3. Results

### 3.1. Estimated Thought States and Their Interpretation

HMM was used to estimate the thought states behind self-reports of thought content and contemplation. The model with four states was employed because the BIC was the smallest (1: 10530.690; 2: 9679.912; 3: 9525.611; 4: 9436.564; 5: 9468.622; 6: 9535.915; 7: 9648.683; 8: 9797.770). The estimated states are shown in Fig. 1A. The vertical axis of the figure indicates thought content or contemplation, and the horizontal axis indicates the probability that each category is selected in each state. In State 1, task-focused had a higher probability of being selected than the other thought contents, but the lowest contemplation was likely to be selected. This is a state in which the participants are focused on the performance of the task and not very thoughtful. In State 2, the probability of selecting task-focused was considerably lower than in State 1, while task-related, interoception, and self-related categories were more prone to be selected. As in State 1, the lowest category of contemplation was more likely to be selected. It is possible that the participants thought vaguely about matters unrelated to the task in this state, and the contents of their thoughts were more likely to be transitional. In State 3, the probability of selecting task-focused was 0%, and task-related or self-related was more likely to be selected. The highest category of contemplation was more likely to be selected. In this condition, the participants rarely focused on the task performance itself but instead thought deeply about things related to the performance of the task or themselves. State 4 was a mind-blanking (MB) condition. Based on the distribution of selected thought content and contemplation, these four thought states were defined as on-task, off-focus, highly contemplative MW, and MB, respectively. Fig. 1B and C show the transition probabilities between the estimated states and the appearance rate of each state, respectively. Among the transitions between the different states, the transition from MB to off-focus had the highest transition probability. On the other hand, transitions from on-task to MB or off-focus and from off-focus to on-task rarely occurred. Transition probabilities between the same state were high for on-task, off-focus, and highly contemplative MW, indicating that these states are more likely to be maintained.

**Fig. 1.**
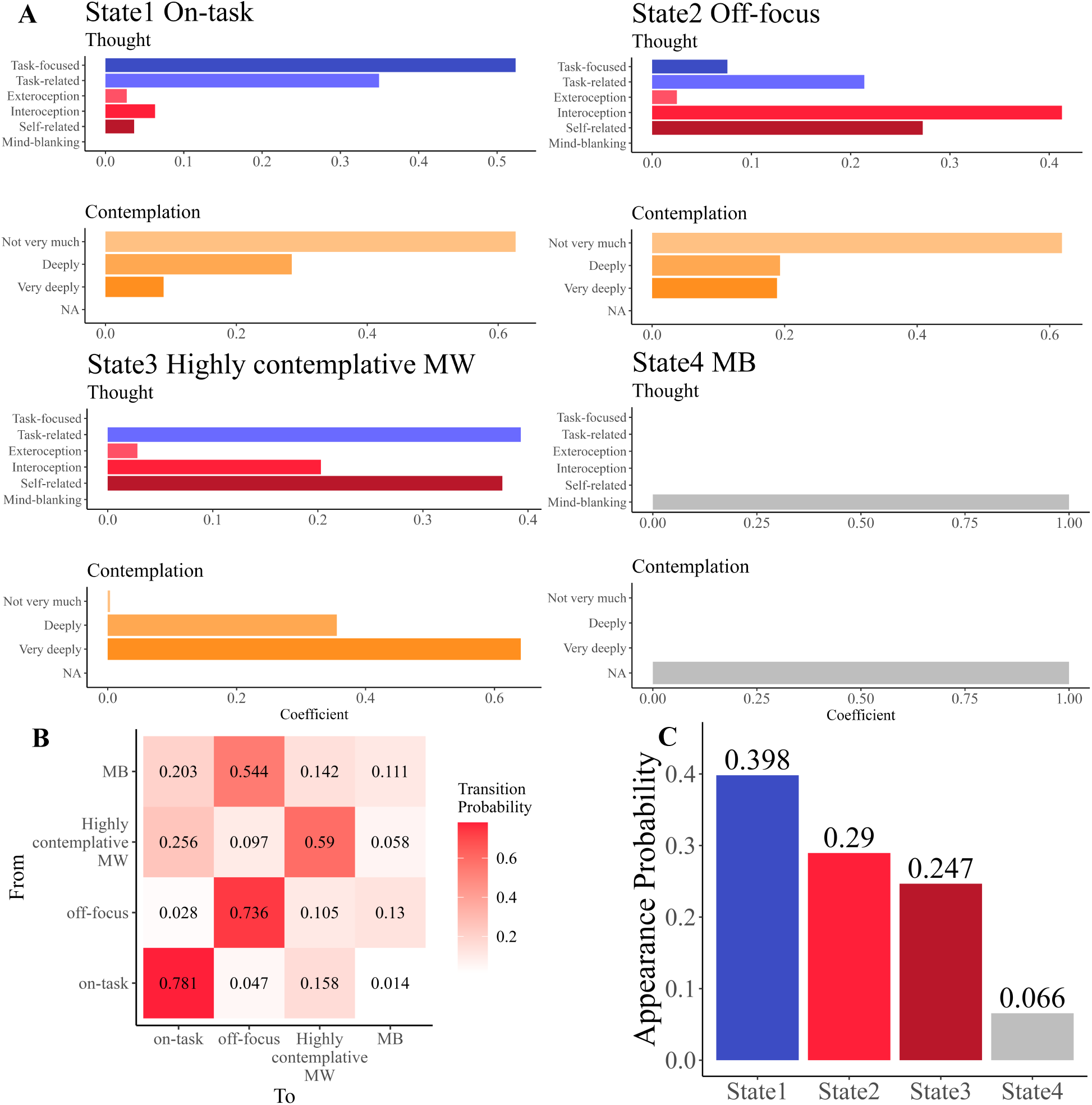
Thought states estimated by HMM. (A) The probability of selecting each thought content and contemplation for each thought state. The vertical axis shows the name of the category of thought content or contemplation, and the horizontal axis shows the probability of selecting each category. Each state was on-task, off-focus, highly contemplative MW, and MB. MW: mind-wandering, MB: mind-blanking. (B) Transition probabilities between thought states. The numbers in the graph indicate the transition probability from the state on the vertical axis to the state on the horizontal axis. The darker the color of the cells in the graph, the higher the transition probability. (C) The appearance probability of each thought state in the task. The horizontal axis indicates the thought state, and the vertical axis indicates the occurrence probability. State1 (on-task) has the highest probability, and State4 (MB) has the lowest probability.

### 3.2. Relationship between the Probability of Thought Transitions During Tasks and Autonomic Nervous Activity

First, to examine whether there were differences in autonomic nervous activity for each of the States indicated in 3.1, we calculated the mean RR and RMSSD for each State. We performed a one-way ANOVA with the State as a factor. No main effect of State was found for either measure (RR: *F* (3) = .602, *p* = .613; RMSSD: *F* (3) = .433, *p* = .729). Thus, there were no differences in autonomic nervous activity by thought states (Appendix Fig. A1, A2). In addition, we also examined whether interoceptive accuracy made a difference in the number of times a particular thought state was reported. The results showed that there was no significant correlation between the number of reports and interoceptive accuracy for any thought states (Fig. A3; State1: *r* = .043, *p* = .773; State2: *r* = .047, *p* = .773; State3: *r* = −.041, *p* = .773; State4: *r* = −.133, *p* = .773). These results indicate that each single thought state is not accompanied by different autonomic nervous activity and that differences in the number of times each thought state is reported are not solely explained by interoceptive accuracy. Based on these results, we decided to examine the relationship between patterns of thought state transitions, autonomic nervous activity, and interoceptive accuracy.

Next, we examined the relationship between the probability of transition to different thought states between successive trials during the task, autonomic nervous activity, and participants’ interoceptive accuracy. We estimated linear regression models with participants’ LF/HF or HF, and the HCT and the TET performances as independent variables and transition probabilities as dependent variables. In addition, we also estimated and calculated the WAIC for models that removed the HCT scores from these models and examined whether including interoceptive accuracy in the models would better explain thought transitions. For the model not including the HCT scores, the WAIC for the LF/HF version was −68.458, and the WAIC for the HF version was −68.949. For the model, including the HCT scores, the WAIC for the LF/HF version was −67.642, and the WAIC for the HF version was −69.251. After comparing the WAIC of these four models, we adopted the HF version model, which included the HCT scores with the lowest WAIC. Information on each parameter in the model is shown in the Appendix (Table A.1-A.3). The results of the analysis of the model included HF, the HCT and the TET performances showed that the interaction between HF and the score of the HCT observed *b* = .336 95% CI [.002;672] (Table 1, Fig.2A). Fig. 2B shows the relationship between the posterior predictive value of transition probability, HF, and the score of the HCT. Fig. 2B and the sign of the interaction coefficient indicate that participants with higher HF during the task and more accurate interoception tended to transition to different thought states.

**Table 1.** Summary of the estimated parameters in transition probability of thought states.

| Parameters | Estimate | Est.Error | l-95% CI | u-95% CI | $\hat{R}$ | Bulk ESS | Tail ESS |
| --- | --- | --- | --- | --- | --- | --- | --- |
| Intercept | 0.536 | 0.127 | 0.287 | 0.787 | 1 | 50892.804 | 42718.938 |
| HF | -10.806 | 6.475 | -23.590 | 1.753 | 1 | 20932.788 | 26950.186 |
| The score of the HCT | -0.002 | 0.001 | -0.005 | 0.001 | 1 | 23751.979 | 31716.620 |
| The score of the TET | -0.001 | 0.001 | -0.004 | 0.001 | 1 | 71573.114 | 40412.126 |
| HF*the HCT | 0.336 | 0.171 | 0.002 | 0.672 | 1 | 20510.139 | 25978.037 |

Notes:
$\hat{R} = 1$ for all parameters, the model has converged
Estimate: the estimated coefficient of each parameter
Est. Error: the standard error in the estimated coefficient of each parameter
l-95% CI: the lower limit of the 95% credible interval (CI)
u-95% CI: the upper limit of the 95% CI
$\hat{R}$ : indicators for determining convergence of Markov Chain Monte Carlo (MCMC) chain
Usually, the model is considered to have converged if $\hat{R} \leq 1.1$
Bulk\_ESS: the adequate sample size calculated by normal rank
Tail\_ESS: the effective sample size for the 5% and 95% quantile points

**Fig. 2.**
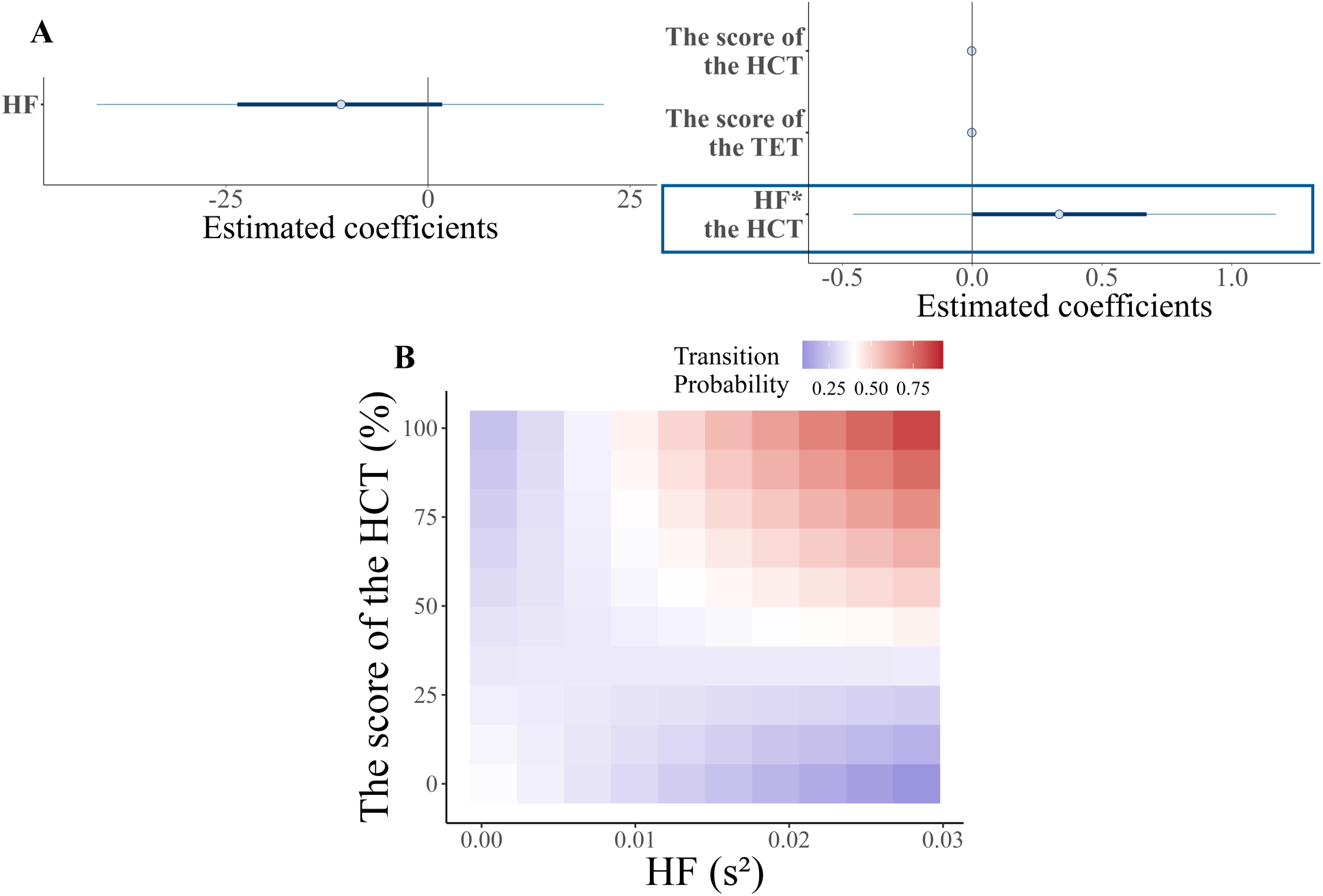
Distribution of the estimated parameters and the posterior prediction of the appearance probability of each transition pattern. (A) The horizontal axis represents the estimated coefficients for each factor. Light blue points indicate the mean of the estimated coefficients, and the dark blue ranges indicate 95% credible intervals (CI). Parameter names in bold boxes indicate that the 95% CI was not across 0. The interaction between HF and interoceptive accuracy influenced the transition probability of the thought states. (B) The vertical axis represents HCT performance, i.e., interoceptive accuracy, and the horizontal axis represents the HF value. The color of the tile represents the transition probability of thought states. The darker the blue, the lower the probability, and the darker the red, the higher the probability. Participants with higher interoceptive accuracy and higher HF during the task were more likely to experience thought transition.

### 3.3. Appearance Probability of Specific Thought Transition Patterns and Trial-by-Trial Autonomic Nervous Activity

We estimated models in which the appearance probabilities of 16 different thought transition patterns are influenced by the interaction between autonomic nervous activity on the trial in which the transition occurs and interoceptive accuracy. Linear mixed models, including individual random effects, were estimated with RMSSD and the score of the HCT as independent variables and the appearance probability of each transition pattern as the dependent variable. We found the interaction between RMSSD and interoceptive accuracy for the transition from State2 (off-focus) to State4 (MB) and the continuation of State3 (highly contemplative MW) (State2→ State4: *b* = .077 95% CI [.018;.135]; State3→State3: *b* = −.065 95%CI [−.121;-.008]; Table 2 and 3, Fig. 3A-1 and 3B-1). Fig. 3A-2 and Fig. 3B-2 show the relationship between the posterior predictive value of appearance probability, RMSSD, and the score of the HCT. From these figures and the signs of the estimated interaction coefficients, participants with more accurate interoception and increased RMSSD on the subsequent trial after off-focus were more likely to move into MB, and participants with more accurate interoception and decreased RMSSD after highly contemplative MW were more likely to continue to experience same thought sate. There was no interaction between RMSSD and interoceptive accuracy for the other transition patterns. For the State2 →State4 and State3→State3 models, we also constructed a model excluding the effect of the HCT scores and estimated the WAIC. The results showed that for both models, the model including the HCT scores better explained the appearance probability of each transition pattern (State2→State4: −60.529; State2→State4 without the HCT: −51.142; State3→State3: −95.490; State3→State3without the HCT: −71.465). Parameters for models other than those in this text are in the Appendix (Appendix Tables A.4-A.19).

**Table 2.** Summary of the estimated parameters in appearance probability of State2→State4.

| Parameters | Estimate | Est.Error | l-95% CI | u-95% CI | $\hat{R}$ | Bulk ESS | Tail ESS |
| --- | --- | --- | --- | --- | --- | --- | --- |
| Intercept | 0.273 | 0.029 | 0.215 | 0.325 | 1 | 36.167 | 21.571 |
| RMSSD | -0.013 | 0.024 | -0.063 | 0.031 | 1 | 234.273 | 8193.895 |
| The score of the HCT | 0.000 | 0.025 | -0.050 | 0.049 | 1 | 130.688 | 13046.775 |
| The score of the TET | -0.015 | 0.022 | -0.059 | 0.028 | 1 | 1375.660 | 18192.256 |
| RMSSD*the HCT | 0.079 | 0.029 | 0.021 | 0.138 | 1 | 1503.429 | 5056.300 |

Notes:
$\hat{R} = 1$ for all parameters, the model has converged
Estimate: the estimated coefficient of each parameter
Est. Error: the standard error in the estimated coefficient of each parameter
l-95% CI: the lower limit of the 95% credible interval (CI)
u-95% CI: the upper limit of the 95% CI
$\hat{R}$ : indicators for determining convergence of Markov Chain Monte Carlo (MCMC) chain
Usually, the model is considered to have converged if $\hat{R} \leq 1.1$
Bulk\_ESS: the adequate sample size calculated by normal rank
Tail\_ESS: the adequate sample size for the 5% and 95% quantile points

**Table 3.** Summary of the estimated parameters in appearance probability of State3→State3.

| Parameters | Estimate | Est.Error | l-95% CI | u-95% CI | $\hat{R}$ | Bulk ESS | Tail ESS |
| --- | --- | --- | --- | --- | --- | --- | --- |
| Intercept | 0.658 | 0.020 | 0.623 | 0.700 | 1 | 82.929 | 473.483 |
| RMSSD | -0.019 | 0.023 | -0.063 | 0.027 | 1 | 107.284 | 444.197 |
| The score of the HCT | 0.009 | 0.019 | -0.029 | 0.043 | 1 | 48.555 | 4771.258 |
| The score of the TET | 0.027 | 0.019 | -0.011 | 0.067 | 1 | 1832.979 | 7485.208 |
| RMSSD*the HCT | -0.075 | 0.026 | -0.128 | -0.022 | 1 | 2261.783 | 2480.872 |

Notes:
$\hat{R} = 1$ for all parameters, the model has converged
Estimate: the estimated coefficient of each parameter
Est. Error: the standard error in the estimated coefficient of each parameter
l-95% CI: the lower limit of the 95% credible interval (CI)
u-95% CI: the upper limit of the 95% CI
$\hat{R}$ : indicators for determining convergence of Markov Chain Monte Carlo (MCMC) chain
Usually, the model is considered to have converged if $\hat{R} \leq 1.1$
Bulk\_ESS: the adequate sample size calculated by normal rank
Tail\_ESS: the adequate sample size for the 5% and 95% quantile points

**Fig. 3.**
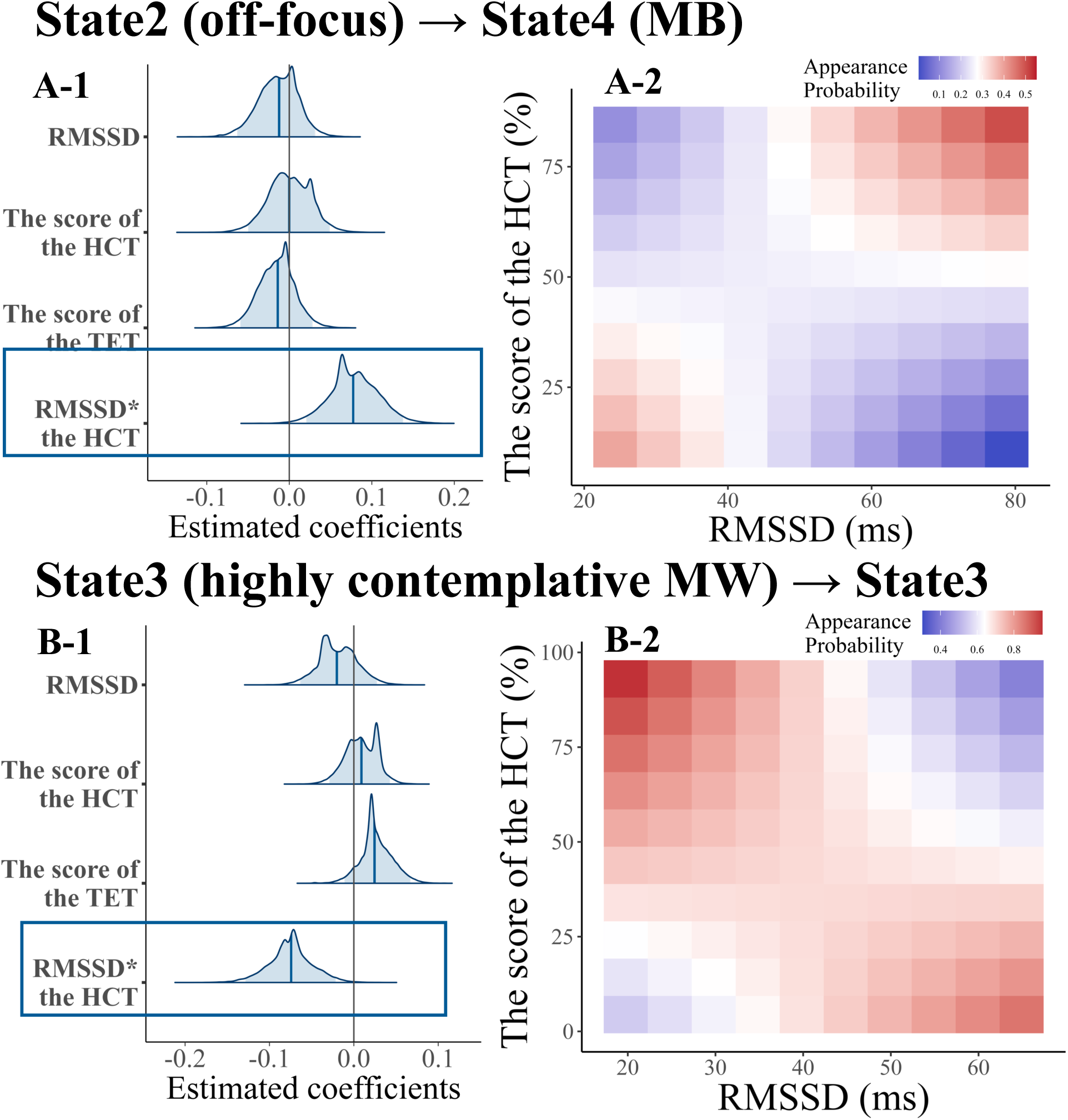
Distribution of the estimated parameters and the posterior prediction of the appearance probability of each transition pattern. (A-1, B-1) The horizontal axis represents the estimated coefficients for each factor. Dark blue lines indicate the mean of the estimated coefficients, and the light blue ranges indicate 95% credible intervals (CI). Parameter names in bold boxes indicate that the 95% CI was not across 0. The interaction between RMSSD and interoceptive accuracy influenced the appearance probability of State2→State4 (A-1) and State3 →State3 (B-1). (A-2, B-2) The vertical axis represents HCT performance, i.e., interoceptive accuracy, and the horizontal axis represents the RMSSD value. The color of the tile represents the appearance probability of the transition pattern of each thought. The darker the blue, the lower the probability; the darker the red, the higher the probability. Participants with higher interoceptive accuracy and higher RMSSD on intertrial were more likely to experience a State2 to State4 transition (A-2). In comparison, participants with higher interoceptive accuracy and lower RMSSD on intertrial were more likely to remain in State 3 (B-2).

## 4. Discussion

The objectives of this study were to estimate the thought state by integrating thought content and contemplation and to examine the relationship between the estimated thought state transitions, fluctuations in autonomic nervous activity, and interoception. Participants were asked to periodically report the content and contemplation of their immediate previous thoughts while participating in a simple attention task. We used the HMM to estimate the latent thought states for time-series data on participants’ thought content and contemplation. The relationship between transition probabilities and transition patterns of thought states, autonomic nervous activity during the task, and interoceptive accuracy was examined. The following two hypotheses were formulated for this study: (1) when parasympathetic activity is dominant, thought state transitions are more frequent, and less contemplative thought states are more likely to occur; conversely, when sympathetic activity is dominant, thought state transitions are less frequent and highly contemplative thought states are more likely to occur, and (2) individual differences in (1) are explained by interoceptive accuracy.

The results of the HMM revealed that the participants’ thoughts during the task consisted of four states (Fig. 1A): focused on task (State. 1; on-task), task-unrelated and low contemplation (State 2; off-focus), task-unrelated and high contemplation (State 3; highly contemplated MW), and no specific thought content reported (State 4; MB). We investigated the relationships among transition patterns in these four distinct thought states, autonomic nervous activity, and interoceptive accuracy. Our findings indicate that participants with higher interoceptive accuracy are more likely to transition among various thought states or to experience shifts between diffuse thought states that lack top-down intentional control (from State 2: off-focus to State 4: MB) when parasympathetic activity is dominant (as indicated by higher values of HF and RMSSD, see Fig. 2B and Fig. 3A-2). Conversely, when sympathetic activity predominated (indicated by low RMSSD values), a propensity for contemplative thought states (State 3: highly contemplated MW) to persist was observed (see Fig. 3B-2). Although these results did not reveal a group-level trend in the correspondence between autonomic nervous activity and thought patterns, as in hypothesis (1), they did reveal a role for an individual’s interoceptive accuracy in mediating the relationship between thought and autonomic nervous activity, as in hypothesis (2).

An interesting finding from this study is that even minute fluctuations in physical responses in the resting state can affect thought states, and that the accuracy of interoception can modulate the interaction between these bodily responses and thought states. In our previous study, subjects with more accurate interoception were more likely to continue engaging in contemplative self-referential thoughts when there were more significant than usual fluctuations in heart rate (Sakuragi et al., 2023). Both these phenomena may be consistent with the concept of the Neural Subjective Frame, which proposes that continuous updating of information about visceral states constructs first-person conscious experiences (Park & Tallon-Baudry, 2014; Tallon-Baudry et al., 2018). Ascending transmission from visceral organs can be divided into two main types: parasympathetic or vagal afferents via cervical ganglia projecting to the solitary bundle nucleus of the brainstem, and sympathetic afferents via dorsal root ganglia projecting through the spinal cord to the brain (Craig, 2002). These signals of autonomic nervous activity are either transmitted to the cortex (Park & Blanke, 2019) or processed in subcortical structures and medial nucleus of the hypothalamus, and later projected to higher brain regions such as the hypothalamus, insular cortex, anterior cingulate cortex, and somatosensory cortex (Critchley & Harrison, 2013). The neural processing of autonomic nervous activity in these higher brain regions shapes and constrains the dynamics of the brain neural network that underpins changes in the states of consciousness (Candia-Rivera, 2022; Menon, 2011; Menon & Uddin, 2010; Uddin, 2015). Among the brain regions associated with information processing in autonomic nervous activity, the insular cortex is also associated with interoceptive accuracy, as measured by the HCT used in this study (Chong et al., 2017). Integrating these previous studies with our results, individuals with higher interoceptive accuracy may be more likely to have more accurate monitoring of their visceral states even at rest, resulting in a switch in brain activity associated with changes in autonomic nervous activity and corresponding transitions in thought states.

On the other hand, the results concerning participants with low interoceptive accuracy should be interpreted with caution. These participants also exhibited different thought transition patterns based on their autonomic nervous activity (Figs. 3A-2, 3B-2). The HCT, employed to assess interoceptive accuracy in this study, measures the accuracy with which individuals can consciously count their heartbeats. However, the actual sensation of pulsation may not always be consciously perceived during the transitions between thought states. Notably, Figs. 3A-2 and 3B-2 illustrate that the relationship between the occurrence of thought transition patterns and autonomic nervous activity of participants with inaccurate interoception is reversed compared to that observed in participants with accurate interoception. This indicates that the mechanisms by which changes in autonomic nervous activity influence thought transitions may vary between individuals with high and low interoceptive accuracy. Future research should comprehensively explore how the link between changes in autonomic nervous activity and thought transitions is mediated by both conscious and unconscious interoception. Employing the Heartbeat Evoked Potential (HEP), which reflects the neural processing of interoception in the unconscious, as a measure, would be particularly beneficial (Park & Blanke, 2019).

This study also revealed the characteristics of the thought transition patterns of people in a resting state. For example, in States 1-3, the continuation of the same thought state is more likely to occur than transitions between different thought states (Fig.1B). This suggests that States 1-3 are more likely to continue, even if there are thought probes presented in between. This result is similar to the trend observed in previous studies that estimated transitional sequences of thought (Shinagawa et al., 2023; Zanesco et al., 2020). In these studies, thought probes were presented on average about once every 20 to 40 seconds; therefore, transitions to different thought states were found to be a relatively rare phenomenon when sampling thoughts at intervals of this length. In the future, it may be possible to clarify the actual timescale of transitions between different thought states by examining the variation in transition probabilities between thought states when the timing of probe presentation is varied. Among the transitions between different states, transitions from MB to off-focus were more likely to occur, while transitions from on-task to MB or off-focus and from off-focus to on-task rarely occurred. This suggests that MB, in which no specific thought content was reported, was similar to off-focus, a diffuse thought state with low contemplation and diverse thought content. However, on-task, in which participants focused on the task despite low contemplation, might differ from these two states. By using the HMM technique as in the present study, it is also possible to estimate the thought state, including more detailed elements of thought (arousal level, valence, tense, etc.). In future studies, we intend to examine what kind of time-series patterns the transition of thought states, including these elements, and whether individual differences in transition patterns can be explained by characteristics such as the state of autonomic nervous activity and interoceptive accuracy.

This study has two limitations. First, it is unclear whether a causal relationship exists between changes in autonomic activity, interoception, and thought transition. Based on the results of this study, it is predicted that autonomic activity changes due to some cause, and that the neural processing of that change alters the overall state of brain activity, which in turn produces a new state of thought. In the future, we plan to measure each dataset and verify the causal relationships among factors using time-series causal inference methods. Second, we did not address individual differences in tendencies of psychiatric disorders, such as anxiety and depression, which are known to be associated with abnormalities in thought, autonomic nervous activity, and interoception. These psychiatric disorder tendencies may affect the frequency of occurrence of the thought states addressed in this study, especially those that are highly contemplative (Ottaviani et al., 2016; Ottaviani, Shahabi, et al., 2015). These individual differences will also be the subject of future research, and the effects of participants’ characteristics on autonomic nervous system activity and thought states will be further investigated.

## 5. Conclusion

This study found that individuals with more accurate interoception were more likely to experience a diffuse state of thought involving less contemplation and more diverse thought content when parasympathetic activity was dominant. Conversely, they were more likely to continue experiencing a highly contemplative state of thought when sympathetic activity was dominant. This is the first study to experimentally examine that autonomic nervous activity in the resting state mirrors the transition pattern of our thought states and that interoception coordinates the relationship between the two. In the future, the influence of autonomic nervous activity and interoception on the generation of various thought states will be clarified by examining thought states that integrate various factors such as valence, tense, and self-relevance of thought, as well as their relationship with psychiatric disorders such as anxiety and depressive tendencies, in which abnormalities in thought have been reported.

## Funding

This work was supported by JSPS KAKENHI Grant Numbers JP18H05525 and JP20H00108 by the Ministry of Education, Culture, Sports, Science and Technology (MEXT), Japan. The authors received no other funding for the research.

## Ethics approval

The study was approved by the Keio University Research Ethics Committee (No. 210300000), Japan.

## Consent to participate

All participants gave informed consent to participate in the present study, and their rights were protected.

## Availability of data and materials

The datasets analyzed during this study are available from the corresponding author upon reasonable request.

## CRediT authorship contribution statement

**Mai Sakuragi:** Conceptualization, Methodology, Software, Formal analysis, Investigation, Writing - Original Draft; **Kazushi Shinagawa:** Software, Formal analysis, Writing - Review & Editing; **Yuri Terasawa:** Writing - Review & Editing; **Satoshi Umeda:** Writing - Review & Editing, Supervision, Funding acquisition.

## Declaration of Competing Interest

The authors declare that they have no known competing financial interests or personal relationships that could have appeared to influence the work reported in this paper.

## Data availability

Data will be made available on request.

## Acknowledgments

This work was supported by JSPS KAKENHI Grant Numbers JP18H05525 and JP20H00108.

## Notes

### Competing Interest Statement

The authors have declared no competing interest.

