## Supplementary material for "The Body Mirroring Thought: The Relationship Between Thought Transitions and Fluctuations in Autonomic Nervous Activity Mediated by Interoception": Fig. A.1.-A.3., Table A.1.-Table A.19.

**Appendix. Supplementally data**

Fig. A1 and A2 show the distribution of RR and RMSSD by State, respectively. The density plots show the distribution of the data; the box plot shows the minimum and maximum values, the first, second, and third quartiles of the data; and the black points next to the box plot show the mean of the data and the error bar show the standard error.

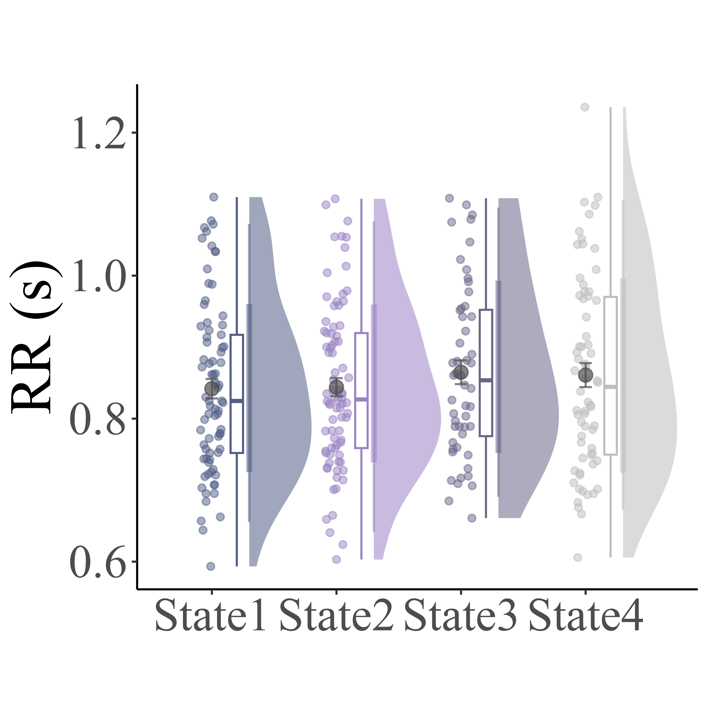

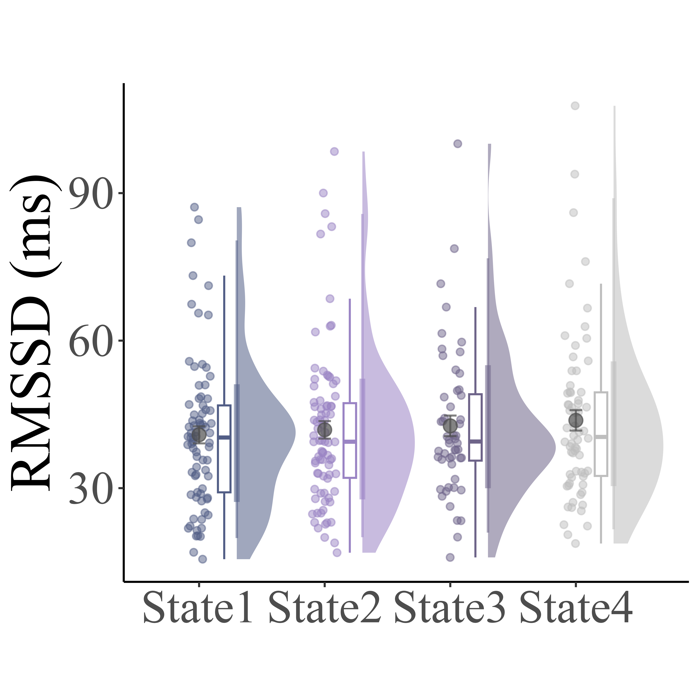

**Fig. A.1. 　　　　　　　　　　　　　　　　　　　Fig. A.2.**

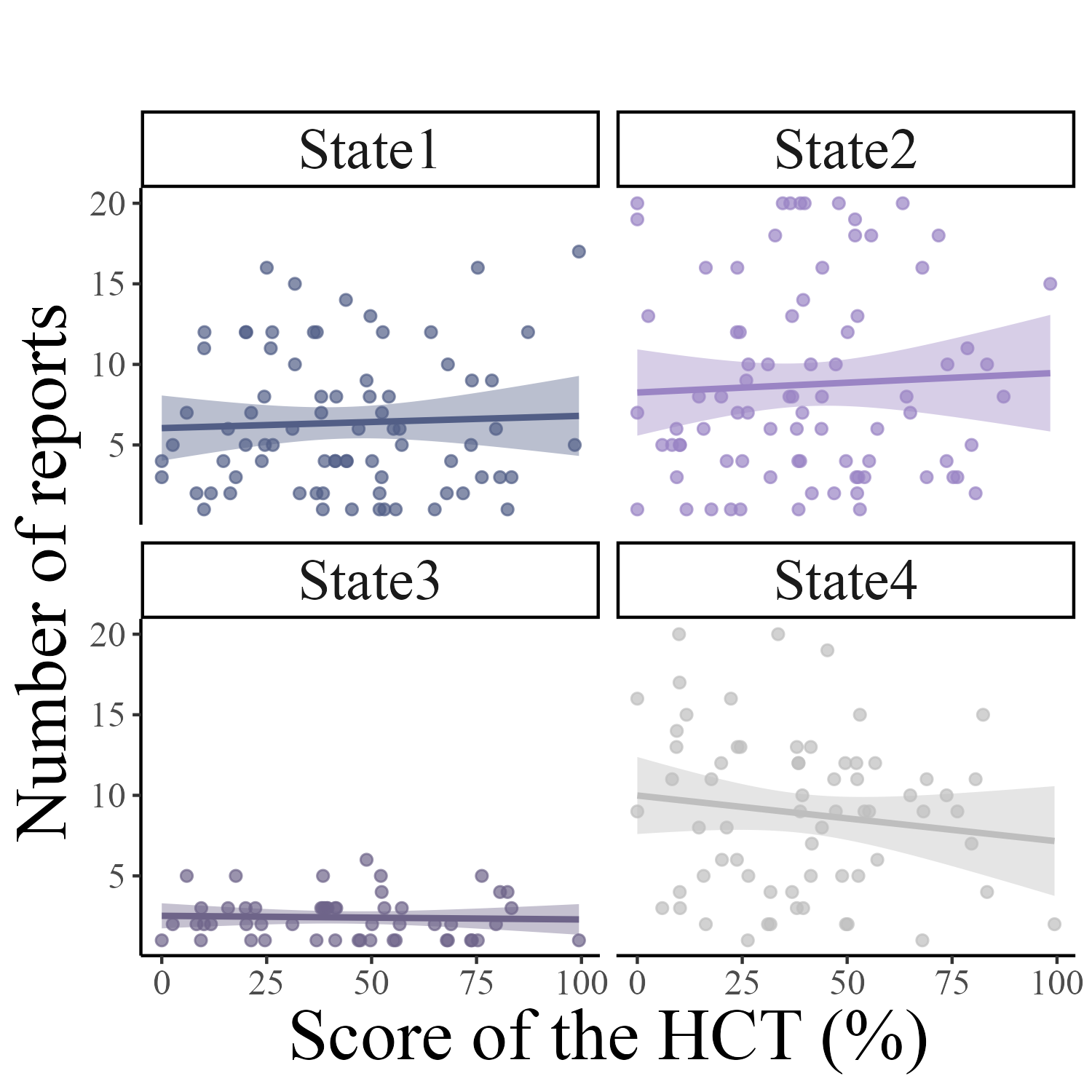

**Fig. A.3.**

**Table A.1.** Summary of the estimated parameters in transition probability of thought states

| Parameters | Estimate | Est.Error | l-95% CI | u-95% CI | Rhat | Bulk_ESS | Tail_ESS |
| --- | --- | --- | --- | --- | --- | --- | --- |
| Intercept | 0.483 | 0.116 | 0.252 | 0.712 | 1 | 69374.909 | 45863.009 |
| LF/HF | 0.000 | 0.005 | -0.010 | 0.009 | 1 | 54525.274 | 38938.071 |
| The score of the TET | -0.001 | 0.001 | -0.004 | 0.001 | 1 | 75092.878 | 43578.261 |

**Table A.2.** Summary of the estimated parameters in transition probability of thought states

| Parameters | Estimate | Est.Error | l-95% CI | u-95% CI | Rhat | Bulk_ESS | Tail_ESS |
| --- | --- | --- | --- | --- | --- | --- | --- |
| Intercept | 0.480 | 0.110 | 0.263 | 0.695 | 1 | 75244.379 | 46726.466 |
| HF | 0.068 | 3.262 | -6.345 | 6.482 | 1 | 50383.721 | 37572.954 |
| The score of the TET | -0.001 | 0.001 | -0.004 | 0.001 | 1 | 78199.386 | 44857.267 |

**Table A.3.** Summary of the estimated parameters in transition probability of thought states

| Parameters | Estimate | Est.Error | l-95% CI | u-95% CI | *R̂* | Bulk_ESS | Tail_ESS |
| --- | --- | --- | --- | --- | --- | --- | --- |
| Intercept | 0.390 | 0.128 | 0.138 | 0.644 | 1 | 47037.961 | 37177.873 |
| LF/HF | 0.014 | 0.010 | -0.005 | 0.033 | 1 | 26923.618 | 27904.140 |
| The score of the HCT | 0.003 | 0.002 | 0.000 | 0.006 | 1 | 33546.692 | 33964.234 |
| The score of the TET | -0.002 | 0.001 | -0.004 | 0.001 | 1 | 58369.477 | 37291.279 |
| LF/HF*the HCT | 0.000 | 0.000 | -0.001 | 0.000 | 1 | 29851.969 | 34030.562 |

**Table A.4.** Summary of the estimated parameters in appearance probability of State1→State1

| Parameters | Estimate | | Est.Error | l-95% CI | u-95% CI | Rhat | Bulk_ESS | Tail_ESS |
| --- | --- | --- | --- | --- | --- | --- | --- | --- |
| Intercept | | 0.732 | 0.020 | 0.694 | 0.771 | 1 | 1535.659 | 3306.914 |
| RMSSD | | 0.009 | 0.020 | -0.032 | 0.047 | 1 | 1303.865 | 611.926 |
| The score of the HCT | | -0.030 | 0.020 | -0.070 | 0.009 | 1 | 2538.634 | 4596.583 |
| The score of the TET | | 0.022 | 0.022 | -0.019 | 0.068 | 1 | 717.458 | 231.702 |
| RMSSD*the HCT | | -0.013 | 0.026 | -0.063 | 0.037 | 1 | 2521.358 | 3148.699 |

**Table A.5.** Summary of the estimated parameters in appearance probability of State1→State2

| Parameters | Estimate | Est.Error | l-95% CI | u-95% CI | Rhat | Bulk_ESS | Tail_ESS |
| --- | --- | --- | --- | --- | --- | --- | --- |
| Intercept | 0.195 | 0.074 | 0.049 | 0.341 | 1 | 25464.683 | 21107.073 |
| RMSSD | -0.004 | 0.050 | -0.103 | 0.095 | 1 | 27442.895 | 30192.063 |
| The score of the HCT | -0.020 | 0.065 | -0.149 | 0.111 | 1 | 27858.350 | 28696.927 |
| The score of the TET | 0.052 | 0.078 | -0.104 | 0.208 | 1 | 24017.732 | 20187.670 |
| RMSSD*the HCT | 0.004 | 0.049 | -0.094 | 0.102 | 1 | 27089.517 | 29295.670 |

**Table A.6.** Summary of the estimated parameters in appearance probability of State1→State3

| Parameters | Estimate | Est.Error | l-95% CI | u-95% CI | Rhat | Bulk_ESS | Tail_ESS |
| --- | --- | --- | --- | --- | --- | --- | --- |
| Intercept | 0.316 | 0.037 | 0.243 | 0.388 | 1 | 4959.666 | 1322.728 |
| RMSSD | 0.016 | 0.043 | -0.067 | 0.100 | 1 | 5426.434 | 17871.906 |
| The score of the HCT | -0.034 | 0.037 | -0.107 | 0.040 | 1 | 12507.324 | 19934.740 |
| The score of the TET | -0.047 | 0.038 | -0.124 | 0.028 | 1 | 6338.271 | 2070.238 |
| RMSSD*the HCT | -0.022 | 0.054 | -0.127 | 0.089 | 1 | 2260.304 | 826.853 |

**Table A.7.** Summary of the estimated parameters in appearance probability of State1→State4

| Parameters | Estimate | Est.Error | l-95% CI | u-95% CI | Rhat | Bulk_ESS | Tail_ESS |
| --- | --- | --- | --- | --- | --- | --- | --- |
| Intercept | 0.059 | 0.665 | -1.380 | 1.465 | 1 | 17796.877 | 11065.962 |
| RMSSD | -0.063 | 0.551 | -1.212 | 1.121 | 1 | 27478.938 | 13595.116 |
| The score of the HCT | 0.002 | 0.510 | -1.073 | 1.065 | 1 | 19665.548 | 11916.773 |
| The score of the TET | -0.073 | 0.501 | -1.155 | 0.977 | 1 | 20240.944 | 12000.742 |
| RMSSD*the HCT | 0.007 | 0.574 | -1.216 | 1.213 | 1 | 25617.850 | 11870.547 |

**Table A.8.** Summary of the estimated parameters in appearance probability of State2→State1

| Parameters | Estimate | Est.Error | l-95% CI | u-95% CI | Rhat | Bulk_ESS | Tail_ESS |
| --- | --- | --- | --- | --- | --- | --- | --- |
| Intercept | 0.210 | 6.982 | -13.601 | 13.882 | 1 | 19289.828 | 9669.404 |
| RMSSD | -0.001 | 3.326 | -6.538 | 6.530 | 1 | 20341.505 | 20737.539 |
| The score of the HCT | -0.027 | 4.985 | -9.779 | 9.781 | 1 | 30130.412 | 12923.817 |
| The score of the TET | -0.031 | 4.825 | -9.471 | 9.427 | 1 | 26385.416 | 16700.774 |
| RMSSD*the HCT | 0.037 | 4.402 | -8.599 | 8.646 | 1 | 27122.763 | 33072.415 |

**Table A.9.** Summary of the estimated parameters in appearance probability of State2→State2

| Parameters | Estimate | Est.Error | l-95% CI | u-95% CI | Rhat | Bulk_ESS | Tail_ESS |
| --- | --- | --- | --- | --- | --- | --- | --- |
| Intercept | 0.714 | 0.021 | 0.673 | 0.755 | 1 | 14297.213 | 19160.681 |
| RMSSD | -0.001 | 0.021 | -0.043 | 0.040 | 1 | 12247.200 | 20500.765 |
| The score of the HCT | -0.020 | 0.022 | -0.063 | 0.023 | 1 | 9707.425 | 9655.209 |
| The score of the TET | 0.016 | 0.019 | -0.022 | 0.054 | 1 | 11533.531 | 17348.227 |
| RMSSD*the HCT | -0.033 | 0.021 | -0.074 | 0.008 | 1 | 8246.680 | 13390.980 |

**Table A.10.** Summary of the estimated parameters in appearance probability of State2→State3

| Parameters | Estimate | Est.Error | l-95% CI | u-95% CI | Rhat | Bulk_ESS | Tail_ESS |
| --- | --- | --- | --- | --- | --- | --- | --- |
| Intercept | 0.224 | 0.057 | 0.110 | 0.336 | 1 | 14394.683 | 19058.569 |
| RMSSD | -0.014 | 0.066 | -0.143 | 0.118 | 1 | 4934.888 | 1833.956 |
| The score of the HCT | -0.051 | 0.055 | -0.159 | 0.054 | 1 | 1106.813 | 1725.957 |
| The score of the TET | 0.023 | 0.052 | -0.079 | 0.126 | 1 | 4858.159 | 24206.417 |
| RMSSD*the HCT | 0.086 | 0.067 | -0.047 | 0.216 | 1 | 2841.803 | 27679.933 |

**Table A.11.** Summary of the estimated parameters in appearance probability of State2→State4

| Parameters | Estimate | Est.Error | l-95% CI | u-95% CI | Rhat | Bulk_ESS | Tail_ESS |
| --- | --- | --- | --- | --- | --- | --- | --- |
| Intercept | 0.256 | 0.026 | 0.204 | 0.306 | 1 | 1133.244 | 9673.308 |
| RMSSD | -0.015 | 0.024 | -0.064 | 0.034 | 1 | 6401.197 | 6077.596 |
| The score of the TET | -0.009 | 0.024 | -0.058 | 0.037 | 1 | 2114.084 | 5366.535 |

**Table A.12.** Summary of the estimated parameters in appearance probability of State3→State1

| Parameters | Estimate | Est.Error | l-95% CI | u-95% CI | Rhat | Bulk_ESS | Tail_ESS |
| --- | --- | --- | --- | --- | --- | --- | --- |
| Intercept | 0.359 | 0.045 | 0.270 | 0.447 | 1 | 8950.066 | 5691.299 |
| RMSSD | 0.037 | 0.047 | -0.055 | 0.129 | 1 | 11161.171 | 14320.981 |
| The score of the HCT | 0.070 | 0.052 | -0.032 | 0.171 | 1 | 4727.257 | 4149.352 |
| The score of the TET | 0.027 | 0.051 | -0.076 | 0.128 | 1 | 7710.184 | 2624.935 |
| RMSSD*the HCT | 0.067 | 0.060 | -0.050 | 0.184 | 1 | 7321.859 | 5037.146 |

**Table A.13.** Summary of the estimated parameters in appearance probability of State3→State2

| Parameters | Estimate | Est.Error | l-95% CI | u-95% CI | Rhat | Bulk_ESS | Tail_ESS |
| --- | --- | --- | --- | --- | --- | --- | --- |
| Intercept | 0.260 | 0.069 | 0.124 | 0.394 | 1 | 7111.983 | 12154.660 |
| RMSSD | 0.004 | 0.094 | -0.181 | 0.191 | 1 | 7797.287 | 13346.361 |
| The score of the HCT | -0.011 | 0.072 | -0.156 | 0.131 | 1 | 5794.333 | 10154.426 |
| The score of the TET | -0.043 | 0.074 | -0.189 | 0.101 | 1 | 6431.107 | 12015.328 |
| RMSSD*the HCT | -0.039 | 0.108 | -0.253 | 0.175 | 1 | 6595.419 | 11699.193 |

**Table A.14.** Summary of the estimated parameters in appearance probability of State3→State3

| Parameters | Estimate | Est.Error | l-95% CI | u-95% CI | Rhat | Bulk_ESS | Tail_ESS |
| --- | --- | --- | --- | --- | --- | --- | --- |
| Intercept | 0.672 | 0.021 | 0.630 | 0.713 | 1 | 11724.622 | 8929.557 |
| RMSSD | -0.010 | 0.024 | -0.057 | 0.039 | 1 | 10477.907 | 13055.383 |
| The score of the TET | 0.017 | 0.021 | -0.024 | 0.058 | 1 | 11244.649 | 15290.649 |

**Table A.15.** Summary of the estimated parameters in appearance probability of State3→State4

| Parameters | Estimate | Est.Error | l-95% CI | u-95% CI | Rhat | Bulk_ESS | Tail_ESS |
| --- | --- | --- | --- | --- | --- | --- | --- |
| Intercept | 0.243 | 0.041 | 0.163 | 0.321 | 1 | 761.900 | 3122.928 |
| RMSSD | 0.017 | 0.060 | -0.103 | 0.135 | 1 | 3327.528 | 3097.391 |
| The score of the HCT | 0.025 | 0.041 | -0.058 | 0.105 | 1 | 5348.849 | 8641.837 |
| The score of the TET | -0.025 | 0.042 | -0.110 | 0.058 | 1 | 1891.370 | 11049.388 |
| RMSSD*the HCT | 0.059 | 0.063 | -0.072 | 0.182 | 1 | 2918.815 | 1477.342 |

**Table A.16.** Summary of the estimated parameters in appearance probability of State4→State1

| Parameters | Estimate | Est.Error | l-95% CI | u-95% CI | Rhat | Bulk_ESS | Tail_ESS |
| --- | --- | --- | --- | --- | --- | --- | --- |
| Intercept | 0.667 | 0.103 | 0.456 | 0.875 | 1 | 12142.189 | 10537.294 |
| RMSSD | 0.155 | 0.110 | -0.065 | 0.371 | 1 | 8966.120 | 4551.949 |
| The score of the HCT | -0.103 | 0.121 | -0.348 | 0.138 | 1 | 12425.656 | 19091.782 |
| The score of the TET | 0.079 | 0.142 | -0.202 | 0.368 | 1 | 6972.081 | 3510.988 |
| RMSSD*the HCT | -0.162 | 0.138 | -0.435 | 0.113 | 1 | 9884.103 | 18294.744 |

**Table A.17.** Summary of the estimated parameters in appearance probability of State4→State2

| Parameters | Estimate | Est.Error | l-95% CI | u-95% CI | Rhat | Bulk_ESS | Tail_ESS |
| --- | --- | --- | --- | --- | --- | --- | --- |
| Intercept | 0.756 | 0.041 | 0.674 | 0.837 | 1 | 19025.705 | 25298.038 |
| RMSSD | -0.011 | 0.038 | -0.087 | 0.064 | 1 | 19518.433 | 26302.004 |
| The score of the HCT | -0.041 | 0.042 | -0.124 | 0.043 | 1 | 18102.572 | 22270.665 |
| The score of the TET | 0.013 | 0.037 | -0.060 | 0.085 | 1 | 17157.984 | 22508.691 |
| RMSSD*the HCT | -0.061 | 0.043 | -0.146 | 0.024 | 1 | 18866.042 | 25181.725 |

**Table A.18.** Summary of the estimated parameters in appearance probability of State4→State3

| Parameters | Estimate | Est.Error | l-95% CI | u-95% CI | Rhat | Bulk_ESS | Tail_ESS |
| --- | --- | --- | --- | --- | --- | --- | --- |
| Intercept | 0.615 | 0.157 | 0.303 | 0.933 | 1 | 29784.188 | 20831.073 |
| RMSSD | -0.011 | 0.155 | -0.320 | 0.300 | 1 | 32219.576 | 28780.803 |
| The score of the HCT | 0.040 | 0.212 | -0.385 | 0.461 | 1 | 24723.131 | 26502.322 |
| The score of the TET | -0.022 | 0.253 | -0.530 | 0.480 | 1 | 26788.298 | 19828.033 |
| RMSSD*the HCT | -0.008 | 0.268 | -0.541 | 0.536 | 1 | 28034.624 | 25431.553 |

**Table A.19.** Summary of the estimated parameters in appearance probability of State4→State4

| Parameters | Estimate | Est.Error | l-95% CI | u-95% CI | Rhat | Bulk_ESS | Tail_ESS |
| --- | --- | --- | --- | --- | --- | --- | --- |
| Intercept | 0.318 | 0.044 | 0.231 | 0.407 | 1 | 29274.168 | 27112.171 |
| RMSSD | 0.039 | 0.048 | -0.058 | 0.133 | 1 | 21292.678 | 18666.978 |
| The score of the HCT | -0.004 | 0.039 | -0.081 | 0.074 | 1 | 29661.718 | 21803.604 |
| The score of the TET | -0.020 | 0.049 | -0.119 | 0.079 | 1 | 27304.143 | 17347.511 |
| RMSSD*the HCT | -0.023 | 0.037 | -0.096 | 0.051 | 1 | 27002.295 | 27868.226 |
